# Bamgineer: Introduction of simulated allele-specific copy number variants into exome and targeted sequence data sets

**DOI:** 10.1101/119636

**Authors:** Soroush Samadian, Jeff P. Bruce, Trevor J. Pugh

## Abstract

Somatic copy number variations (CNVs) play a crucial role in development of many human cancers. The broad availability of next-generation sequencing data has enabled the development of algorithms to computationally infer CNV profiles from a variety of data types including exome and targeted sequence data; currently the most prevalent types of cancer genomics data. However, systemic evaluation and comparison of these tools remains challenging due to a lack of ground truth reference sets. To address this need, we have developed Bamgineer, a tool written in Python to introduce user-defined haplotype-phased allele-specific copy number events into an existing Binary Alignment Mapping (BAM) file, with a focus on targeted and exome sequencing experiments. As input, this tool requires a read alignment file (BAM format), lists of non-overlapping genome coordinates for introduction of gains and losses (bed file), and an optional file defining known haplotypes (vcf format). To improve runtime performance, Bamgineer introduces the desired CNVs in parallel using queuing and parallel processing on a local machine or on a high-performance computing cluster. As proof-of-principle, we applied Bamgineer to a single high-coverage (mean: 220X) exome sequence file from a blood sample to simulate copy number profiles of 3 exemplar tumors from each of 10 tumor types at 5 tumor cellularity levels (20-100%, 150 BAM files in total). To demonstrate feasibility beyond exome data, we introduced read alignments to a targeted 5-gene cell-free DNA sequencing library to simulate *EGFR* amplifications at frequencies consistent with circulating tumor DNA (10, 1, 0.1 and 0.01%) while retaining the multimodal insert size distribution of the original data. We expect Bamgineer to be of use for development and systematic benchmarking of CNV calling algorithms by users using locally-generated data for a variety of applications. The source code is freely available at http://github.com/pughlab/bamgineer.

**Author summary:** We present Bamgineer, a software program to introduce user-defined, haplotype-specific copy number variants (CNVs) at any frequency into standard Binary Alignment Mapping (BAM) files. Copy number gains are simulated by introducing new DNA sequencing read pairs sampled from existing reads and modified to contain SNPs of the haplotype of interest. This approach retains biases of the original data such as local coverage, strand bias, and insert size. Deletions are simulated by removing reads corresponding to one or both haplotypes. In our proof-of-principle study, we simulated copy number profiles from 10 cancer types at varying cellularity levels typically encountered in clinical samples. We also demonstrated introduction of low frequency CNVs into cell-free DNA sequencing data that retained the bimodal fragment size distribution characteristic of these data. Bamgineer is flexible and enables users to simulate CNVs that reflect characteristics of locally-generated sequence files and can be used for many applications including development and benchmarking of CNV inference tools for a variety of data types.

## Introduction

The emergence and maturation of next-generation sequencing technologies, including whole genome sequencing, whole exome sequencing, and targeted sequencing approaches, has enabled researchers to perform increasingly more complex analysis of copy number variants (CNVs)[1]. While genome sequencing-based methods have long been used for CNV detection, these methods can be confounded when applied to exome and targeted sequencing data due to noncontiguous and highly-variable nature of coverage and other biases introduced during enrichment of target regions[1–5]. In cancer, this analysis is further challenged by bulk tumor samples that often yield nucleic acids of variable quality and are composed of a mixture of cell-types, including normal stromal cells, infiltrating immune cells, and subclonal cancer cell populations. Circulating tumor DNA presents further challenges due to a multimodal DNA fragment size distribution and low amounts of tumor-derived DNA in blood plasma. Therefore, development of CNV calling methods on arbitrary sets of tumor-derived data from public repositories may not reflect the type of tumor specimens encountered at an individual centre, particularly formalin-fixed-paraffin embedded tissues routinely profiled for diagnostic testing.

Due to lack of a ground truth for validating CNV callers, many studies have used simulation studies to model tumor data[6]. Most often, simulation studies are used in an *ad-hoc* manner using customized formats to validate specific tools and settings with limited adaptability to other tools. More generalizable approaches aim at the *de novo* generation of sequencing reads according to a reference genome (e.g. wessim[3], Art-illumina[7], and dwgsim[8]. However, *de novo* simulated reads do not necessarily capture subtle features of empiric data, such as read coverage distribution, DNA fragment insert size, quality scores, error rates, strand bias and GC content[6]; factors that can be more variable for exome and targeted sequencing data particularly when derived from clinical specimens. Recently, Ewing et al. developed a tool, BAMSurgeon, to introduce synthetic mutations into existing reads in a Binary alignment Mapping (BAM) file[9]. BAMSurgeon provides support for adjusting variant allele fractions (VAF) of engineered mutations based on prior knowledge of overlapping CNVs but does not currently support direct simulation of CNVs themselves.

Here we introduce Bamgineer, a tool to modify existing BAM files to precisely model allele-specific and haplotype-phased CNVs (Fig 1). This is done by introducing new read pairs sampled from existing reads, thereby retaining biases of the original data such as local coverage, strand bias, and insert size. As input, Bamgineer requires a BAM file and two lists of nonoverlapping genomic coordinates to introduce allele-specific gains and losses. The user may explicitly provide known haplotypes or chose to use the BEAGLE[10] phasing module that we have incorporated within Bamgineer. We implemented parallelization of the Bamgineer algorithm for both standalone and high performance computing cluster environments, significantly improving the scalability of the algorithm. Overall, Bamgineer gives investigators complete control to introduce CNVs of arbitrary size, magnitude, and haplotype into an existing reference BAM file. We have uploaded all software code to a public repository (http://github.com/pughlab/bamgineer).

**Fig 1.**
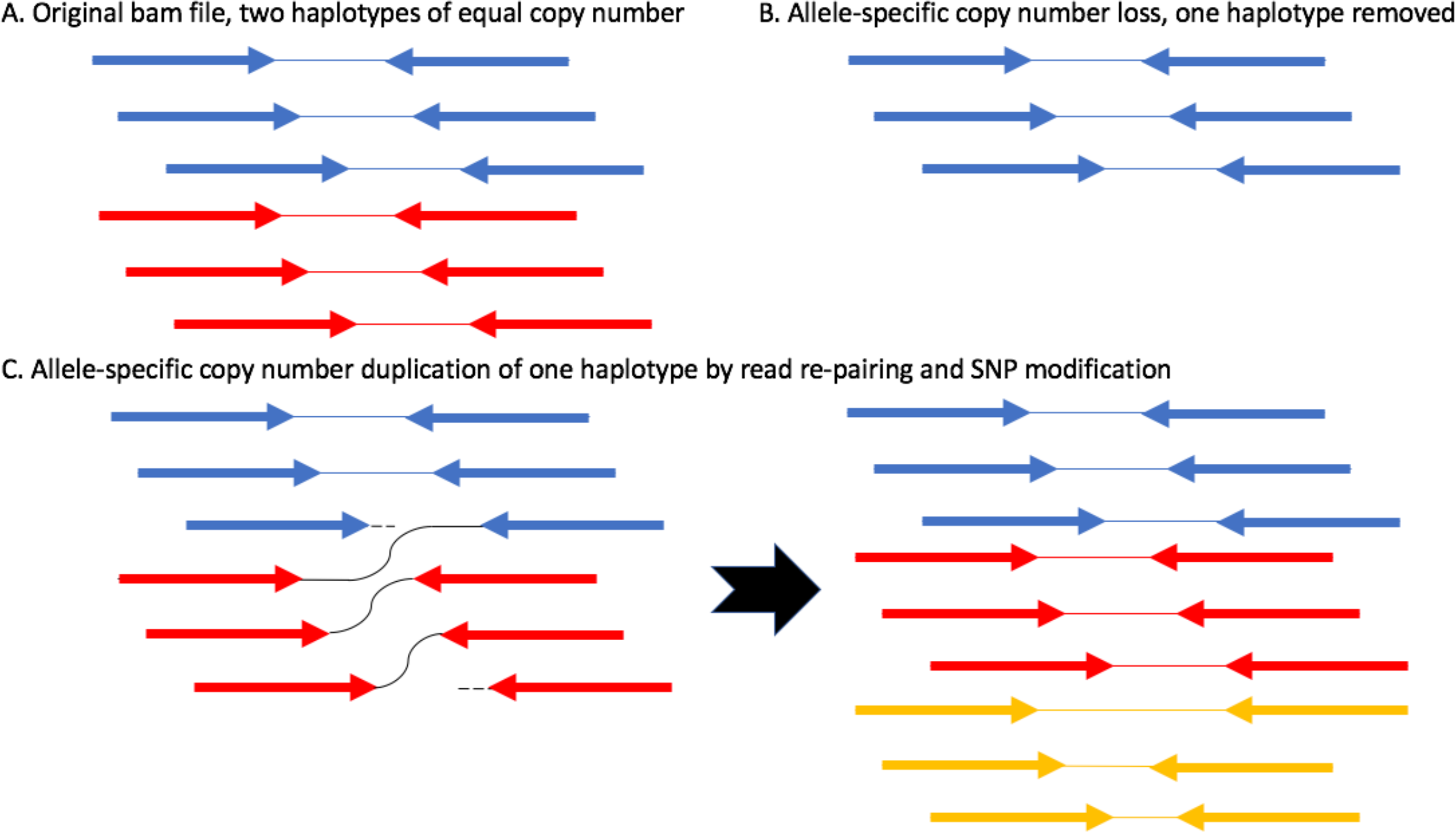
Bamgineer conceptual overview. Haploype-specific CNVs simulated using re-paired reads. Red and blue colors represent read-pairs corresponding to different haplotypes. Orange color represents new reads corresponding to the red haplotype. **A**. Original BAM file used as the input. **B**. Allele-specific loss, reads matching a target haplotype are removed **C**. Allele-specific gain for copy number equal 3. New read pairs are constructed from existing reads and reads are modified at SNP loci to match the desired haplotype.

## Results

### Proof-of-principle experiments using whole exome sequence data

For all proof-of-principle experiments, we used exome sequencing data from a single normal (peripheral blood lymphocyte) DNA sample. DNA was captured using the Agilent SureSelect Exome v5+UTR kit and sequenced to 220X median coverage as part of a study of neuroendocrine tumors. Reads were aligned to the hg19 build of the human genome reference sequence and processed using the Genome Analysis Toolkit (GATK) Best Practices pipeline.

#### Arm-level and chromosome-level copy number alteration

To verify the ability of our workflow to introduce copy gains, we first generated single-copy, allele-specific copy number amplification (ASCNA) of chromosomes 21 and 22, followed by calculation of hybrid-selection metrics using the Picard suite[11]. As expected, the number of read pairs increased by 50% in the two chromosomal regions of interest with no statistically significant impact median, mean, and standard deviation of insert size for paired-reads (Table S1). The distribution of paired reads distances was nearly indistinguishable before and after the addition of ASCNA (median 211.97 vs. 212.3, Two-sided KS-test p>0.99, S5 Fig). We subsequently applied our exome analysis pipeline for detecting CNVs using Mpileup[12], Varscan2[13] and Sequenza[14] to infer allele-specific copy number profiles (S6 Fig). This analysis detected the desired arm- and chromosome-level gains at the expected depth ratio of 1.5 and we verified specific gain of desired haplotypes by confirming that 97-98% of the reads contained variants corresponding to the target haplotype (mean variant allele frequency of SNPs on amplified haplotypes = 0.66±**0. 03** and 0.33 versus 0.50±0.03 on non-target haplotypes; Fig 2A and B).

**Fig 2.**
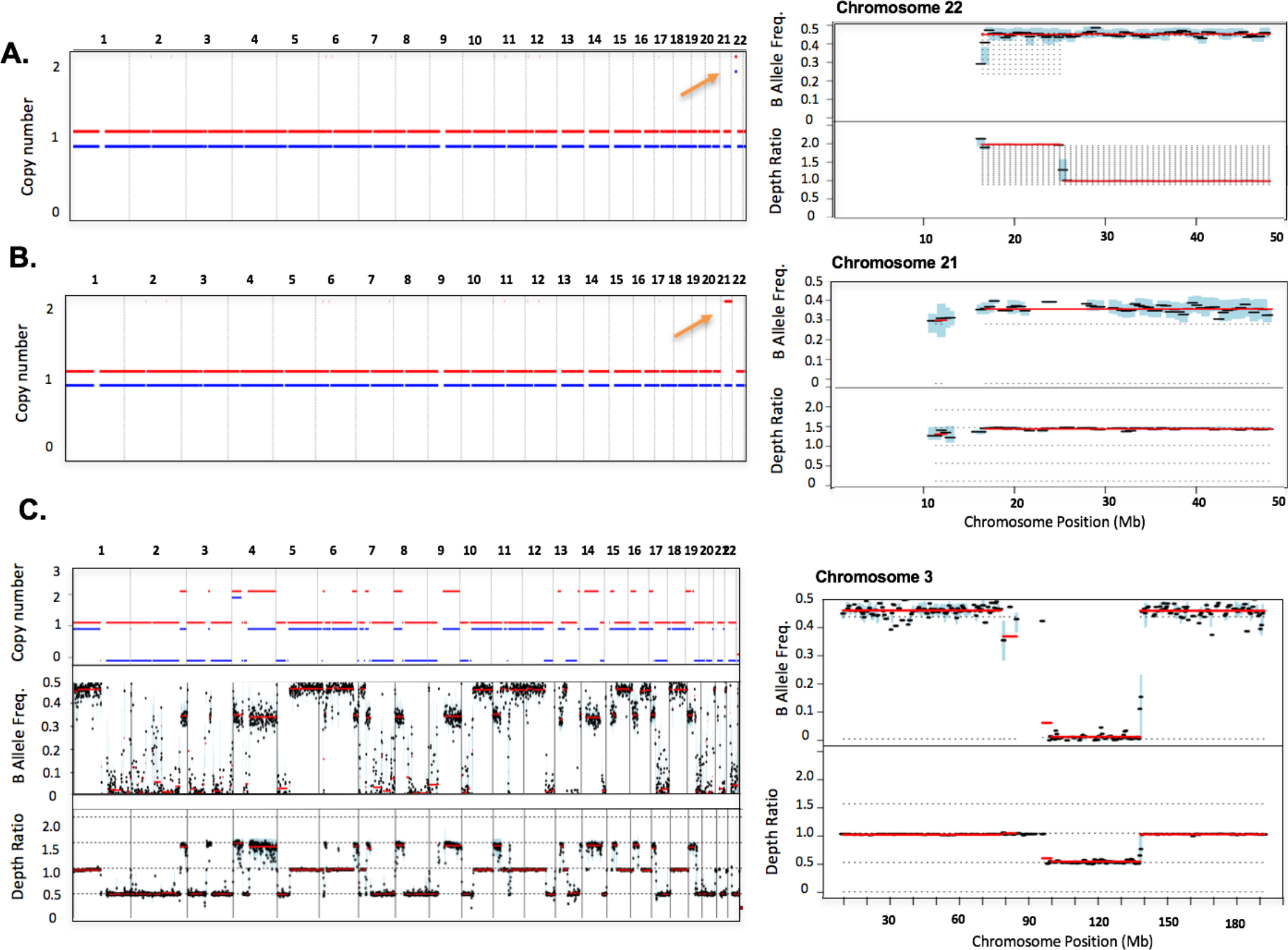
Example of allele specific CNV calls generated from modified bam files. A) Genome-wide (left) and chromosome-view (right) of allele specific copy number, BAF and depth ratios for balanced gain of p-arm in chromosome 22 inferred using Sequenza. Blue and red lines show allele specific copy number profiles for each chromosome (lines are offset from discrete copy number values by ± 0.1 for visual separation of the two alleles). The small blue and red spots on the top figure (orange circle) show a balanced gain on p-arm of chromosome 22 (BAF is not affected as a result of balanced gain). Each black dot on the right figures represents a genomic locus and the red lines indicate the inferred value for consecutive segments. B) Allele-specific gain of entire chromosome 21(orange circle). As shown only one copy of the chromosome is gained and hence the allele frequency is reduced from the 0.5 to ~0.33 in the chromosome view. C) Genome-wide (left) and chromosome-view (right) for 36 events (21 gains and 25 losses) sampled from Genome Atlas for Bladder Urothelial Carcinoma (BLCA) for 100% tumor content. As expected depth ratio and BAFs are approximately 0.5 and zero respectively.

We next repeated this experiment to introduce a single copy loss of these two regions. Again, the number of read pairs was consistent with the desired copy state (decrease of 50% of reads in the target regions). We also verified that the single copy deletion was restricted to a single haplotype with > 99% of reads containing variants corresponding to the target haplotype.

### Synthetic tumor-normal mixtures of exemplar tumors from TCGA

Following the validation of our tool for readily-detected chromosome- and arm-level events, we next used Bamgineer to simulate CNV profiles mimicking 3 exemplar tumors from each of 10 different cancer types profiled by The Cancer Genome Atlas using the Affymetrix SNP6 microarray platform: lung adenocarcinoma (LUAD); lung squamous cell (LUSC); head and neck squamous cell carcinoma (HNSC); glioblastoma multiformae (GBM); kidney renal cell carcinoma (KIRC); bladder (BLCA); colorectal (CRC); uterine cervix (UCEC); ovarian (OV), and breast (BRCA) cancers (Table 1).

**Table 1.** Average number of copy number gains, losses and percent genome altered (PGA) across TCGA cancer types and the three exemplar tumors selected for our study.

| TCGA |  |  | Exemplar tumors |  |  |  |  |  |  |  |  |
| --- | --- | --- | --- | --- | --- | --- | --- | --- | --- | --- | --- |
| Average of all tumors |  |  | Tumor 1 |  |  | Tumor 2 |  |  | Tumor 3 |  |  |
| Cancer | Gains | Losses | Gains | Losses | PGA | Gains | Losses | PGA | Gains | Losses | PGA |
| BLCA | 23 | 30 | 23 | 30 | 70% | 22 | 25 | 72% | 22 | 25 | 71% |
| BRCA | 26 | 36 | 24 | 34 | 59% | 27 | 33 | 67% | 21 | 30 | 66% |
| CRC | 24 | 42 | 27 | 48 | 27% | 23 | 29 | 70% | 22 | 30 | 69% |
| GBM | 26 | 36 | 23 | 34 | 51% | 19 | 33 | 73% | 30 | 43 | 57% |
| HNSC | 24 | 40 | 28 | 40 | 61% | 24 | 30 | 55% | 24 | 28 | 62% |
| KIRC | 6 | 16 | 6 | 16 | 75% | 7 | 15 | 57% | 8 | 14 | 66% |
| LUAD | 26 | 34 | 26 | 25 | 55% | 26 | 30 | 63% | 22 | 32 | 67% |
| LUSC | 25 | 43 | 34 | 47 | 26% | 31 | 34 | 68% | 24 | 31 | 73% |
| UCEC | 39 | 47 | 39 | 43 | 62% | 38 | 38 | 59% | 37 | 37 | 66% |
| OV | 29 | 37 | 26 | 35 | 68% | 28 | 34 | 74% | 28 | 31 | 65% |

To select 3 exemplar tumors for each cancer type, we chose profiles that best represented the copy number landscape for each cancer type. First, we addressed over-segmentation of the CNV calls from the microarray data by merging segments of <500 kb in size with the closest adjacent segment and removing the smaller event from the overlapping gain and loss regions. We then assigned a score to each tumor that reflects its similarity to other tumor of the same cancer type (S7 Fig). This score integrates total number of CNV gain and losses (Methods, Equation 6), median size of each gain and loss, and the overlap of CNV regions with GISTIC peaks for each cancer type as reported by The Cancer Genome Atlas (Table 1). We selected three high ranking tumors for each cancer type such that, together, all significant GISTIC[15] peaks for that tumor type were represented. A representative profile from a single tumor is shown in Fig 2C.

Subsequently, for each of the 30 selected tumor profiles (3 for each of 10 cancer types), we introduced the corresponding CNVs at 5 levels of tumor cellularity (20, 40, 60, 80, and 100%) resulting in 150 BAM files in total. For each BAM file, we used Sequenza to generate allele-specific copy number calls as done previously. Tumor/normal log2 ratios are shown in Fig 3 for one representative from each cancer type. From this large set of tumors, we next set out to compare Picard metrics and CNV calls as we did for the arm- and chromosome-level pilot.

**Fig 3.**
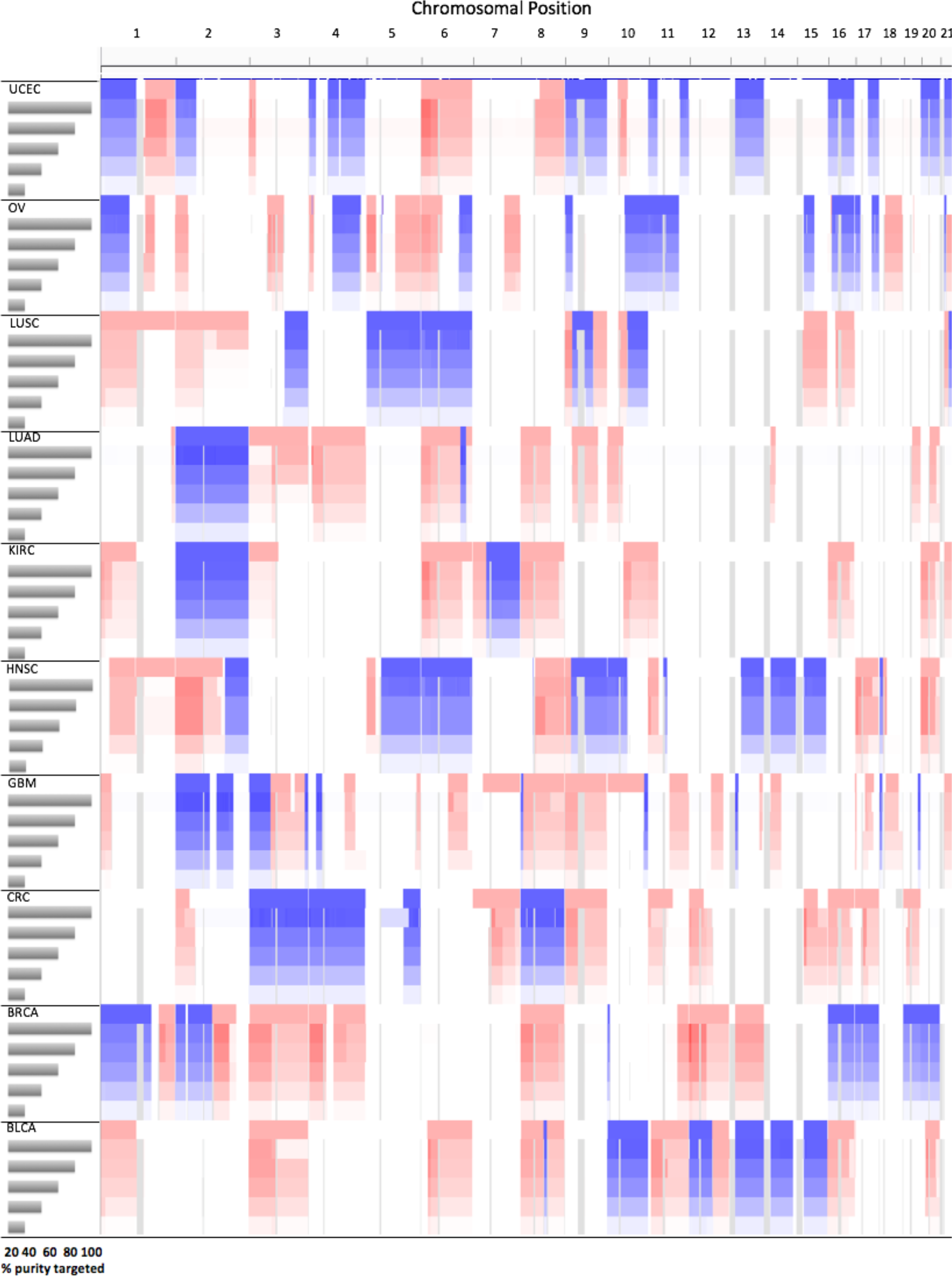
Log2 ratios from simulated exemplar tumors at varying purity levels. Log2 ratios of copy number segments inferred using the Sequenza algorithm, shown as a heatmap (blue: loss, red: gain; data range is −1.5 to 1.5) for different cancers and different tumor cellularities shown for one representative tumor for each cancer type. The representative tumor is created by combining high-ranking tumor profiles from TCGA segments for each cancer (defined by similarity score; Equation 6). The quantized segments from representative tumor is used as the CNV input for the exemplar tumors. A single normal BAM file is used as the input file to generate each representative tumor at 100% purity; subsequently, 100% tumor and the original normal are sampled at (100%, 80%, 60%, 40% and 20%) to artificially create tumor-normal admixture at the desired purity. The purity value for each tumor type (in gray), is the median purity value for each cancer type according to TCGA segments. As shown in the figure, as the purity decreases, we observe a corresponding decrease in tumor segments (reduction in color intensities).

### Performance evaluation

We evaluated Bamgineer using several metrics: tumor allelic ratio, SNP phasing consistency, and tumor to normal log2 ratios (Fig 4). As expected, across all regions of a single copy gain, tumor allelic ratio was at ~0.66 (interquartile range: 0.62 - 0.7) for the targeted haplotype and 0.33 (interquartile range: 0.3-0.36) for the other haplotype. As purity was decreased, we observed a corresponding decrease in allelic ratios, from 0.66 down to 0.54 (interquartile range: 0.5 - 0.57) for targeted and an increase (from 0.33) to 0.47 (interquartile range: 0.43-0.5) for the other haplotype for 20% purity (Fig 4A and B). These changes correlated directly with decreasing purity (R^2^ > 0.99) for both haplotypes. Similarly, for single copy loss regions, as purity was decreased from 100% to 20% the allelic ratio linearly decreased (R^2^ > 0.99) from ~0.99 (interquartile range: 0.98-1.0) for targeted haplotype to ~0.55 (interquartile range: 0.51-0.58) for targeted haplotype and increases from 0 to ~0.43 (interquartile range: 0.4-0.46) for the other haplotype (Fig 4B). The results for log2 tumor to normal depth ratios of segments normalized for average ploidy were also consistent with the expected values (Methods, Equation 2). For CNV gain regions, log2 ratio decreased from ~0.58 (log2 of 3/2) to ~0.13 as purity was decreased from 100% to 20%. For CNV loss regions, as purity was decreased from 100% to 20%, the log2 ratio increased from −1 (log2 of 1/2) to −0.15, consistent with Equation 2 (Fig 4C; S1-S4 for individual cancers).

**Fig 4.**
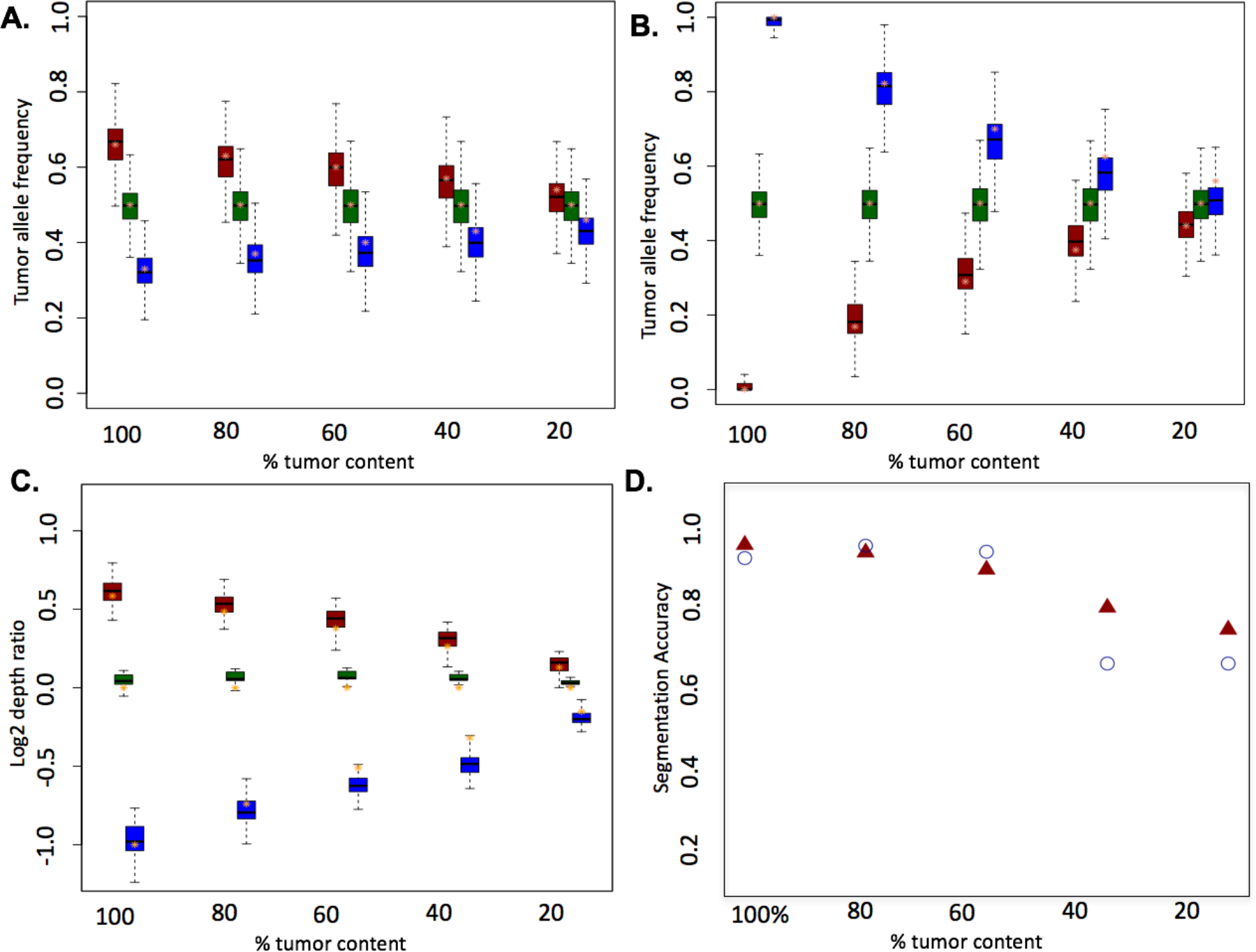
Exome-wide simulated copy number profiles at a range of tumor purities yield expected allelic and copy number ratios. Allelic ratio for allele-specific copy number gain (A) and loss (B) events at heterozygous SNP loci for haplotypes affected (blue), haplotypes not affected (red), and SNPs not in engineered CNV regions (green) as negative controls at different tumor cellularity levels (x-axis) across all cancers. C) Tumor to normal log2 depth ratio boxplots of copy number gain (red) and loss (blue) segments from Sequenza across all cancers (Table 1). D) Accuracy of Sequenza copy segment calling gains (red triangles) and losses (blue circles) decreases as simulated tumor content decreases.

Ultimately, we wanted to assess whether Bamgineer was introducing callable CNVs consistent with segments corresponding to the exemplar tumor set. To assess this, we calculated an accuracy metric (Fig 4B) as:

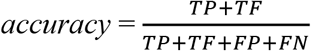

where TP, TF, FP and FN represent number of calls from Sequenza corresponding to true positives (perfect matches to desired CNVs), true negatives (regions without CNVs introduced), false positives (CNV calls outside of target regions) and false negatives (target regions without CNVs called). TP, TF, TN, FN were calculated by comparing Sequenza absolute copy number (predicted) to the target regions for introduction of 1 Mb CNV bins across the genome. As tumor content decreased, accuracy for both gains and losses decreased as false negatives became increasingly prevalent due to small shifts in log2 ratios. We note that (as expected), overall decreasing cancer purity from 100% to 20% generally decreases the segmentation accuracy. Additionally, we observe that segmentation accuracy is on average, significantly higher for gain regions compared to the loss regions for tumor purity levels below 40% (Fig 4D). This is consistent with previous studies that show the sensitivity of CNV detection from sequencing data is slightly higher for CNV gains compared to CNV losses[16]. We also note that with decreasing cancer purity, the decline in segmentation accuracy follows a linear pattern of decline for gain regions and an abrupt stepwise decline for loss regions (Fig 4D; segmentation accuracies are approximately similar for 40% and 20% tumor purities).

Finally, we observed a degree of variation in terms of segmentation accuracy across individual cancer types (S1-S4 Fig). Segmentation accuracy was lower for LUAD, OV and UCEC compared to other simulated cancer types for this study. The relative decline in performance is seen in cancer types where CNV gains and losses cover a sizeable portion of the genome; and hence, the original loss and gain events sampled from TCGA had significant overlaps. As a result, after resolving overlapping gain and loss regions (S7 Fig), on average, the final target regions constitute a larger number of small (< 200 kb) loss regions immediately followed by gain regions and vice versa; making the accurate segmentation challenging for the CBS (circular binary segmentation) algorithm implemented by Sequenza relying on presence of heterozygous SNPs. This, can cause uncertainties in assignments of segment boundaries.

In summary, application of an allele-specific caller to BAMs generated by Bamgineer recapitulated CNV segments consistent with >95% (medians: 95.1 for losses and 97.2 for gains) of those input to the algorithm. However, we note some discrepancies between the expected and called events, primarily due to small CVNs as well as large segments of unprobed genome between exonic sequences.

### Synthetic tumor-normal mixtures using cell free DNA sequence data

To evaluate the use of Bamgineer for circulating tumor DNA analysis, we simulated the presence of an *EGFR* gene amplification in read alignments from a targeted 5-gene panel (18 kb) applied to a cell-free DNA from a healthy donor and sequenced to >50,000X coverage. To mirror concentrations of tumor-derived fragments commonly encountered in cell-free DNA, we introduced gain of an *EGFR* haplotype at frequencies of 100, 10, 1, 0.1, and 0.01%. This haplotype included 3 SNPs covered by our panel, which were phased and subject to allele-specific gain accordingly. As with the exome data, we observed shifts in coverage of specific allelic variants, and haplotype representation consistent with the targeted allele frequencies (Fig 5A, Supplemental Table S1). Furthermore, read pairs introduced to simulate gene amplification retain the bimodal insert size distribution characteristic of cell-free DNA fragments (Fig 5B and C).

**Fig 5.**
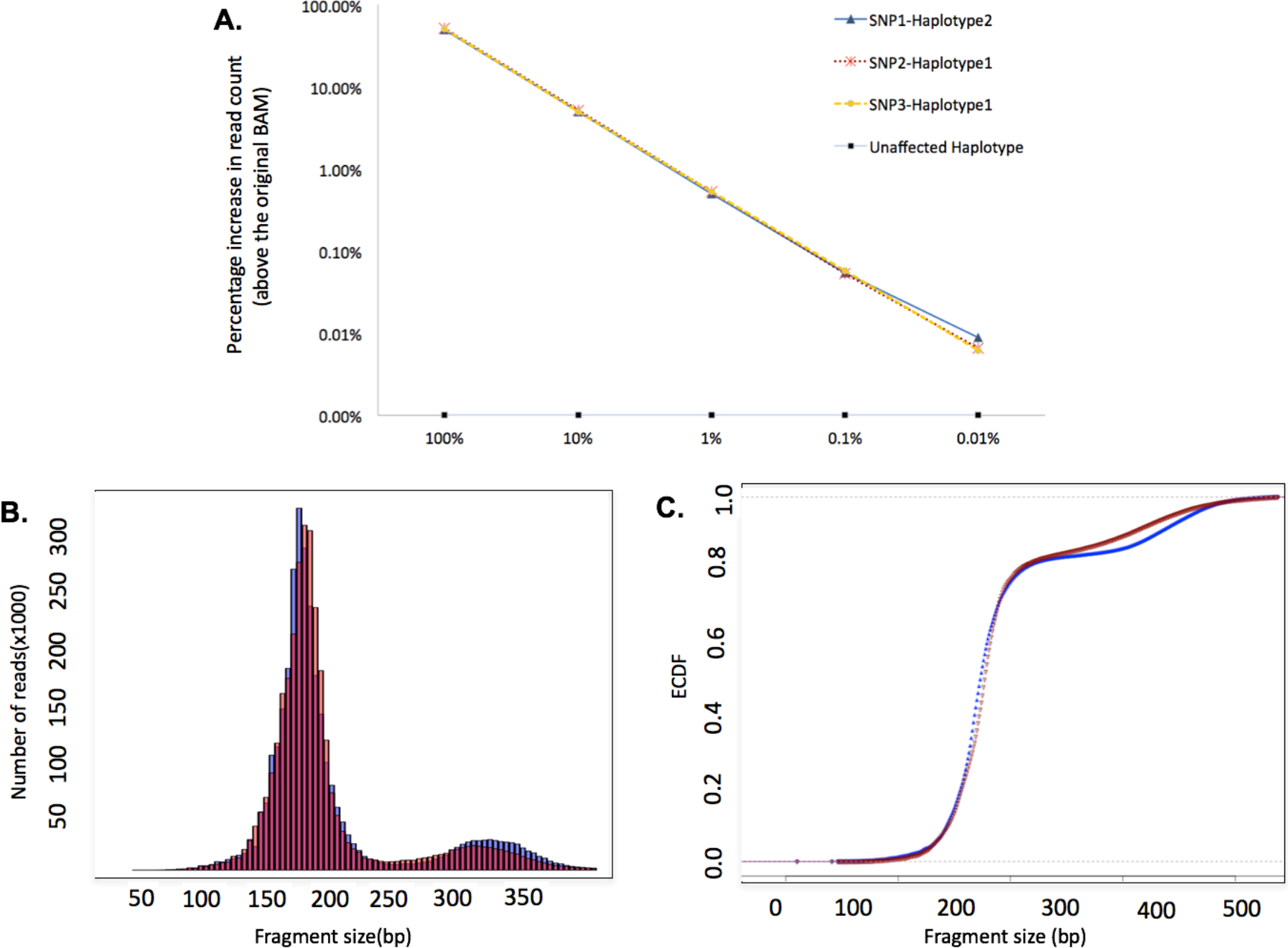
Simulated low frequency CNVs in circulating tumor DNA data yield expected allelic ratios and retain underlying bimodal fragment size distribution. A) Percentage increase in read count above the original input BAM file for the affected haplotype for three SNPs in EGFR region. Shifts in coverage of specific allelic variants, and haplotype representation consistent with the targeted allele frequencies. B) Comparison of DNA inserts size distribution from a targeted 5-gene cell-free DNA sequencing library subject to introduction of read pairs supporting an *EGFR* single-copy gain. Distribution of insert sizes using 5bp bin size. C) Experimental Cumulative Density Functions (ECDF) of all fragment lengths (blue; median 173, mean: 194.5) and newly introduced read pairs (red; median 175, mean: 197.7) allele specific gain for EGFR. Despite the multimodal nature of the cell free DNA distribution of fragment size (two major peaks at ~160 and 330bp), the fragment size distribution of the original read pairs and that of read pairs introduced to simulate EGFR gain are reasonably consistent (Two sided KS test: 0.11: p-value: 0.81; we note minor discrepancies in relative intensity of second peak at around ~330 relative to the original BAM).

While this experiment showcases the ability of Bamgineer to faithfully represent features of original sequencing data while controlling allelic amplification at the level of the individual reads, these subtle shifts are currently beyond the sensitivity of conventional CNV callers when applied to small, deeply covered gene panels. Therefore, it is our hope that Bamgineer may be of value to aid develop of new methods capable of detecting copy number variants supported by a small minority of DNA fragments in a specimen.

### Runtime and parallelization

Bamgineer is computationally intensive and the runtime of the program is dictated by the number of reads that must be processed, a function of the coverage of the genomic footprint of target regions. To ameliorate the computational intensiveness of the algorithm, we employed a parallelized computing framework to maximize use of a high-performance compute cluster environment when available. We took advantage of two features in designing the parallelization module. First, we required that added CNVs are independent for each chromosome (although nested events can likely be engineered through serial application of Bamgineer). Second, since we did not model interchromosomal CNV events, each chromosome can be processed independently. As such, CNV regions for each chromosome can be processed in parallel and aggregated as a final step. S8 Fig shows the runtimes for The Cancer Genome Atlas simulation experiments. Using a single node with 12 cores and 128 GB of RAM, each synthetic BAM took less than 3.5 hours to generate. We also developed a version of Bamgineer that can be launched from sun grid engine cluster environments. It uses python pipeline management package *ruffus* to parallelize tasks automatically and log runtime events. It is highly modular and easily updatable. If disrupted during a run, the pipeline can continue to completion without re-running previously completed intermediate steps.

## Discussion

Here, we introduced Bamgineer, to introduce user-defined haplotype-phased allele-specific copy number events into an existing Binary Alignment Mapping (BAM) file, obtained from exome and targeted sequencing experiments. As proof of principle, we generated, from a single high coverage (mean: 220X) BAM file derived from a human blood sample, a series of 30 new BAM files containing a total of 1,693 simulated copy number variants (on average, 56 CNVs comprising 1800Mb i.e. ~55% of the genome per tumor) corresponding to profiles from exemplar tumors for each of 10 cancer types. To demonstrate quantitative introduction of CNVs, we further simulated 4 levels of tumor cellularity (20, 40, 60, 80% purity) resulting in an additional 120 new tumor BAM files. We validated our approach by comparing CNV calls and inferred purity values generated by an allele-specific CNV-caller (Sequenza[14]) as well as a focused comparison of allelic variant ratios, haplotype-phasing consistency, and tumor/normal log2 ratios for inferred CNV segments (S1-S4 Fig). In every case, inferred purity values were within ±5% of the targeted purity; and majority of engineered CNV regions were correctly called by Sequenza (accuracy > 94%; S1-S4 Fig). Allele variant ratios were also consistent with the expected values both for targeted and the other haplotypes (Median within ±3% of expected value). Median tumor/normal log2 ratios were within ±5% of the expected values.

To demonstrate feasibility beyond exome data, we next evaluated these same metrics in a targeted 5-gene panel applied to a cell-free DNA sequencing library generated from a healthy blood donor and sequenced to >10,000X coverage[17] To simulate concentrations of tumor-derived fragments typically encountered in cancer patients, we introduced *EGFR* amplifications at frequencies of 100, 10, 1, 0.1, and 0.01%. As with the exome data, we observed highly specific shifts in allele variant ratios, log2 coverage ratios, and haplotype representation consistent with the targeted allele frequencies. Our method also retained the bimodal DNA insert size distribution observed in the original read alignment. However, it is worthwhile noting that, these minute shifts are currently beyond the sensitivity of existing CNV callers when applied to small, deeply covered gene panels. Consequently, we anticipate that Bamgineer may be of value to aid develop of new methods capable of detecting copy number variants supported by a small minority of DNA fragments.

The significance of this work in the context of CNV inference in cancer is twofold: 1) users can simulate CNVs using their own locally-generated alignments so as to reflect lab-, biospecimen-, or pipeline-specific features; 2) bioinformatic methods development can be better supported by ground-truth sequencing data reflecting CNVs without reliance on generated test data from suboptimal tissue or plasma specimens. Bamgineer addresses both problems by creating standardized sequencing alignment format (BAM files) harbouring user-defined CNVs that can readily be used for algorithm optimization, benchmarking and other purposes.

We expect our approach to be applicable to tune algorithms for detection of subtle CNV signals such as somatic mosaicism or circulating tumor DNA. As these subtle shifts are beyond the sensitivity of many CNV callers, we expect our tool to be of value for the development of new methods for detecting such events trained on conventional DNA sequencing data. By providing the ability to create customized user-generated reference data, Bamgineer will prove valuable inn development and benchmarking of CNV calling and other sequence data analysis tools and pipelines.

The work presented herein can be extended in several directions. First, Bamgineer is not able to reliably perform interchromosomal operations such as chromosomal translocation, as our focus has been on discrete regions probed by exome and targeted panels. Additionally, in our current implementation, we limited the maximum total copy number to 4. Certainly, higher-level amplifications occur in cancer and iterative application of Bamgineer may enable introduction of copy number states beyond four chromosomal copies as well as complex, nested events. While these are challenging events to model, we appreciate that this class of structural variant play an important driving role in cancer; for example, EGFR vIII variants observed in brain cancer. Finally, introduction of compound, serially acquired CNVs may be of interest to model subclonal phylogeny developed over time in bulk tumor tissue samples.

## Materials and Methods

Bamgineer uses several python packages for parsing input files (pyVCF[18], VCFtools[19], and pybedtools[20], manipulating BAM files (pysam[21]; Samtools[22], Sambamba[23] and BamUtils[24]), phasing haplotypes (BEAGLE[10], and distributing compute jobs in cluster environments (ruffus[25]). HaplotypeCaller[26] from the Genome Analysis Toolkit (GATK) is used to call germline heterozygous SNPs (het.vcf) if known haplotype SNP data is not provided. The analysis workflow is outlined in Fig 6.

**Fig 6.**
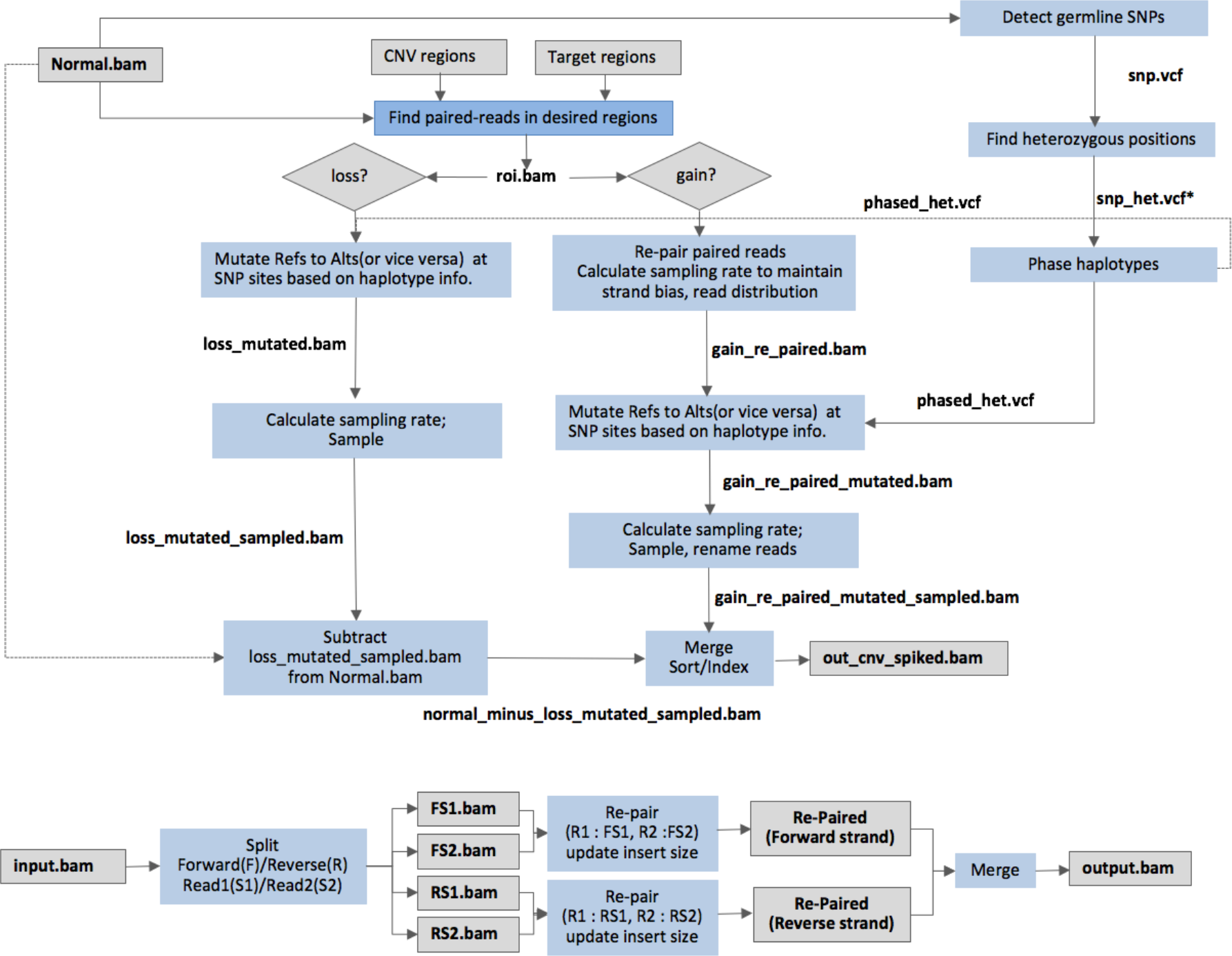
Bamgineer algorithm overview. A) Overall architecture of Bamgineer for editing an existing BAM file to add and delete the user defined CNV event. The input and output files are shown in dark grey. The modules are shown darker relative to the files generated at each step. B) Creating new paired-reads from existing ones. The algorithms splits the input BAM file into four separate files according to DNA strand and read information. Bamgineer then iterates through split reads (read1 and read2) from each strand separately, pairing one read from read1 splits to another read from read2 split. The insert-size (tlen) in the newly paired read is then calculated and updated.

### Inputs

The user provides 2 mandatory inputs to Bamgineer as command-line arguments: 1) a BAM file containing aligned paired-end sequencing reads (“*Normal.bam*”), 2) a BED file containing the genome coordinates and type of CNV (e.g. allele-specific gain) to introduce (“CNV regions.bed”). Bamgineer can be used to add four broad categories of CNVs: Balanced Copy Number Gain (BCNG), Allele-specific Copy Number Gain (ASCNG), Allele-specific Copy Number Loss (ACNL), and Homozygous Deletion (HD). For example, consider a genotype AB at a genomic locus where A represents the major and B represents the minor allele.

Bamgineer can be applied to convert that genomic locus to any of the following copy number states:

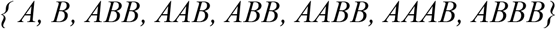

An optional VCF file containing phased germline calls can be provided (*phased_het.vcf*). If this file is not provided, Bamgineer will call germline heterozygous single nucleotide polymorphisms (SNPs) using the GATK HaplotypeCaller and then categorize alleles likely to be co-located on the same haplotypes using BEAGLE and population reference data from the HapMap project.

### Isolation of source reads to construct haplotype-specific CNVs

To obtain paired-reads in CNV regions of interest, we first intersect *Normal.bam* with the targeted regions overlapping user-defined CNV regions (*roi.bed*). This operation generates a new BAM file (*roi.bam*). Subsequently, depending on whether the CNV event is a gain or loss, the algorithms performs two separate steps as follows.

### I. Creation of new read pairs simulating copy gains

To introduce copy number gains, Bamgineer creates new read-pairs constructed from existing reads within each region of interest. This approach thereby avoids introducing pairs that many tools would flag as molecular duplicates due to read 1 and read 2 having start and end positions identical to an existing pair. If desired, these read pairs can be restricted to reads meeting a specific SAM flag. For our exome experiments, we used read pairs with a SAM flag equal to 99, 147, 83, or 163, i.e. read paired, read mapped in proper pair, mate reverse (forward) strand, and first (second) in pair. To enable support for the bimodal distribution of DNA fragment sizes in ctDNA, we removed the requirement for “read mapped in proper pair” and used read pairs with a SAM flag equal to 97, 145, 81, or 161. Users considering engineering of reads supporting large inserts or intrachromosomal read pairs may also want to remove the requirement for “read mapped in proper pair”. Additionally, we required that the selection of the newly paired read is within ±50%(±20% for ctDNA) of the original read size.

The newly created read- pairs are provided unique read names to avoid confusion with the original input BAM file. To enable inspection of these reads, these newly created read pairs are stored in a new BAM file, *gain_re_paired_renamed.bam*, prior to merging into the final engineered BAM. Since we only consider high quality reads (i.e. properly paired reads, primary alignments and mapping quality > 30), the newly created BAM file contains fewer reads compared to the input file (~90-95% in our proof-of-principle experiment). As such, at every transition we log the ratio between number of reads between the input and output files.

#### Introduction of mutations according to haplotype state

To ensure newly constructed read-pairs match the desired haplotype, we alter the base at heterozygous SNP locations (*phasedhet.vcf*) within each read according to haplotype provided by the user or inferred using the BEAGLE algorithm. To achieve this, we iterate through the set of re-paired reads used to increase coverage (*gain_re_paired_renamed.bam*) and modify bases overlapping SNPs corresponding to the target haplotype (*phased_het.vcf*). We then write these reads to a new BAM file (*gain_re_paired_renamed_mutated.bam*) prior to merging into the final engineered BAM (Fig 6).

As an illustrative example consider two heterozygous SNPs, *AB* and *CD* both with allele frequencies of ~0.5 in the original BAM file (i.e. approximately half of the reads supporting reference bases and the other half supporting alternate bases. To introduce a 2-copy gain of a single haplotype, reads to be introduced must match the desired haplotype rather than the two haplotypes found in the original data. If heterozygous *AB* and *CD* are both located on a haplotype comprised of alternative alleles, at the end of this step, 100% of the newly re-paired reads will support alternate base-pairs (e.g. BB and DD). Based on the haplotype structure provided, other haplotype combinations are possible including AA/DD, BB/CC, etc.

#### Sampling of reads to reflect desired allele fraction

Depending on the absolute copy number desired for the for CNV gain regions, we sample the BAM files according to the desired copy number state. We define conversion coefficient as the ratio of total reads in the created BAM from previous step (*gain_repaired_mutated.bam*) to the total reads extracted from original input file (*roi.bam*):

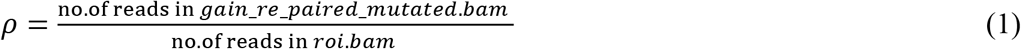

According to the maximum number of absolute copy number (ACN) for simulated CNV gain regions (defined by the user), two scenarios are conceivable as follows.

#### Single copy gain (ACNG = 3)

To achieve the single copy gain (ACN =3, e.g. ABB copy state), the file in the previous step (*gain_re_paired_renamed_mutated.bam*), should be subsampled such that on average depth of coverage is half that of extracted reads from the target regions from the original input normal file(*roi.bam*). Thus, the final sampling rate is calculated by dividing 0.5 by *ρ* (subsample *gain_re_paired_renamed_mutated.bam* such that we have half of the *roi*.bam depth of coverage for the region; in practice adjusted sampling rate is in the range of 0.51-0.59 i.e. 0.85 < *ρ* < 1) and the new reads are written to a new BAM file (*gain_re_paired_renamed_mutated_sampled.bam*) that we then merge with the original reads *(roi.bam)* to obtain *gain_final.bam*.

#### Double copy gain (ACNG = 4)

To achieve a 2-copy gain (ACN =4, e.g. AABB copy state), the average depth of coverage for input file *roi.bam* and sampled BAM file, should be the equal. However since *ρ* < 1, we subsample of the input reads (*roi.bam*) such that on average depth of coverage of the sampled file (*roi_sampled.bam*) is equal that of the synthetic BAM file (*gain_re_paired_renamed_mutated.bam*). We subsequently merge the two files to generate *gain_final.bam*. Note that this effectively will generate a BAM file with slightly (0.85 < *ρ* < 1) lower depth of coverage than the original file.

### II. Removing read-pairs to simulate CNV Loss

To introduce CNV losses, Bamgineer removes reads from the original bam corresponding to a specific haplotype and does not create new read pairs from existing ones. To diminish coverage in regions of simulated copy number loss, we sub-sample the BAM files according to the desired copy number state and write these to a new file. The conversion coefficient is defined similarly as the number of reads in *rai_loss_mutated.bam* divided by number of reads in *roiloss.bam* (> ~0.98). Similar to CNV gains, the sampling rate is adjusted such that after the sampling, the average depth of coverage is half that of extracted reads from the target regions (calculated by dividing 0.5 by conversion ratio, as the absolute copy number is 1 for loss regions). Finally, we subtract the reads in CNV loss BAMs from the *input.bam* (or *input_sampled.bam*) and merge the results with CNV gain BAM (*gain_final.bam*) to obtain, the final output BAM file harbouring the desired copy number events.

### Statistical benchmarking and evaluation

To validate that the new paired-reads generated from the original BAM files show similar probability distribution, we used two-sided Kolmogorov-Smirnov (KS) test. The critical D-values where calculated for *α* = 0.01 as follows:

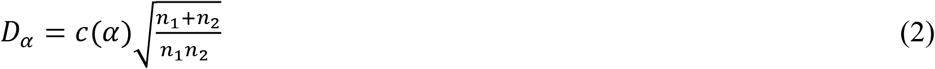

where coefficient c(α) is obtained from Table of critical values for KS test (https://www.webdepot.umontreal.ca/Usagers/angers/MonDepotPublic/STT3500H10/Critical_KS.pdf; 1.63 for α = 0.01) and n_1_ and n_2_ are the number of samples in each dataset. To assess tumor allelic ratio consistency, for each SNP the theoretical allele frequency parameter was used as a reference point (Equation 3). Median, interquartile range and mean were drawn from the observed values for each haplotype-event pair for all the SNPs. The boxplot distribution of the allele frequencies were plotted and compared against the theoretical reference point. To assess the segmentation accuracy, we used log2 tumor to normal depth ratios of segments normalized for mean ploidy as the metric; where the mean ploidy is (Equations 4 and 5). To benchmark the performance of segmentation accuracy, we used accuracy as the metrics. Statistical analysis was performed with the functions in the R statistical computing package using RStudio.

#### Theoretical expected values

The expected value for tumor allelic frequencies at heterozygous SNP *loci* for tumor purity level of *p* (1-*p*: normal contamination) is calculated as follows:

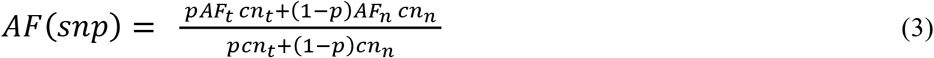

where *AF*_*t*_ and *AF*_*n*_ represent the expected allele frequencies for tumor and normal and *cn*_*t*_ and *cn*_*n*_ the expected copy number for tumor and normal at specific SNP *loci*. For CNV events used in this experiment *AF*_*t*_ are (1/3 or 2/3) for gain and (1 or 0) for loss CNVs according to the haplotype information (whether or not they are located on the haplotype that is affected by each CNV). The expected value for the average ploidy 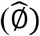 is calculated as follows

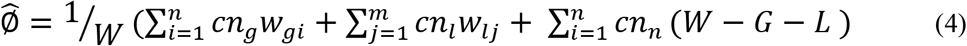

where *cn*_*g*_, *cn*_*l*_, *cn*_*n*_, *w*_*g*_ and *w*_*l*_ represent the expected ploidy for gain, loss and normal regions, and the length of individual gain and loss events respectively. *W, G, and L* represent total length (in base pairs) for gain regions, loss regions, and the entire genome (~ 3e9).

The expected log2ratio for each segment is calculated as follows

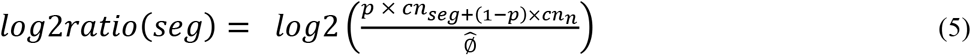

where *cn*_*seg*_ is the segment mean from Sequenza output, *p* tumor purity and 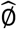 is the average ploidy calculated above. *cn*_*n*_ is the copy number of copy neutral region (i.e. 2)

#### Similarity score to rank TCGA tumors

The similarity score for specific cancer type (c) and sampled tumor (t) is calculated as follows:

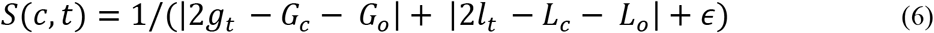

where *g*_*t*_, *G*_*c*_, *G*_*o*_ represent the total number of gains for specific tumor sampled from Cancer Genome Atlas (after merging adjacent regions and removing overlapping regions), median number of gains for specific tumor type, and number of gain events overlapping with GISTIC peaks respectively; *l*_*t*_, *L*_*c*_, *L*_*o*_ represent the above quantities for CNV loss regions(*ϵ* is an arbitrary small positive value to avoid zero denominator.). The higher score the closer is the sampled tumor to an exemplar tumor from specific cancer type.

## Acknowledgements

We thank Rene Quevedo, Olena Kis, the staff of the Princess Margaret Genomics Centre (www.pmgenomics.ca, Neil Winegarden, Julissa Tsao and Nick Khuu) and Bioinformatics Services (Carl Virtanen, Zhibin Lu and Natalie Stickle) for their expertise in generating the sequencing data used in this study. This work was supported by the Princess Margaret Cancer Foundation (T.J.P.), Canada Foundation for Innovation, Leaders Opportunity Fund, CFI #32383 (T.J.P.); Ontario Ministry of Research and Innovation, Ontario Research Fund Small Infrastructure Program (T.J.P.).

## Supporting information captions

**S1 Fig. Cancer-specific allelic ratio (y-axis) changes with purity (x-axis) for CNV gain regions.** Allelic ratio boxplots for cancer-specific copy number gain at heterozygous SNP *loci* for haplotypes affected (blue) and Haplotypes not affected (red) at different tumour cellularity levels.

**S2 Fig. Cancer-specific allelic ratio (y-axis) changes with purity (x-axis) for CNV loss regions.** Allelic ratio boxplots for cancer-specific copy number loss at heterozygous SNP *loci* for haplotypes affected (blue) and Haplotypes not affected (red) at different tumour cellularity levels.

**S3 Fig. Cancer-specific tumor/normal log2 ratio(y-axis) changes with purity(x-axis).** Tumor to normal log2 depth ratio boxplots for cancer-specific copy number gain (red) and loss (blue) at different tumour cellularity levels normalized for mean ploidy.

**S4 Fig. Cancer-specific segmentation accuracy changes with purity.**

**S5 Fig. ECDFs of original and newly created reads.** Experimental Cumulative Density Functions (ECDF) of all fragment lengths (blue; median 194, mean: 212.3) and newly introduced read pairs (red; median 194, mean: 211.97) allele specific gain for chromosome 22. The distribution before and after the addition of CNV is consistent.

**S6 Fig. Whole Exome Sequencing pipeline for CNV inference.** Pipeline for detecting absolute and allele-specific CNV. Varscan2 was used for depth normalization. Varscan2 output was then used to infer allele-specific copy number profiles as well as tumour purity and ploidy using Sequenza note that in this case “tumour” file will be the synthetic file generated using Bamgineer.

**S7 Fig. Cancer-specific CNV introduction.** Overview of the design used to introduce cancer-specific CNV events. Parallelization module enables to simultaneously implement cancer-based, chromosome-based and event-based engineering of CNVs, significantly improving the performance (see “Runtime benchmarks and parallelization”)

**S8 Fig. Bamgineer runtimes (hours) for different tumour types.**

**S1 Table.** Allele counts at 3 SNPS in *EGFR*. Introduction of *EGFR* gain to targeted 5-gene panel (18 kb) applied to a cell-free DNA at frequencies of 100, 10, 1, 0.1, and 0.01%. The Alt and Ref columns represent the count of alternative and reference base pairs at each variant position. The columns in bold represents the phased targeted haplotype. We note that Bamgineer can be used to introduce subtle shifts in coverage of specific allelic variants, and haplotype representation consistent with the targeted allele frequencies.

| Tumour Purity | Alt | Ref | Exp.(Alt) | Exp.(Ref) |
| --- | --- | --- | --- | --- |
| SNP1 (rs2227983) |  |  |  |  |
| Original BAM | 32260 | **31769** | 32260 | **31769** |
| 100% | 32260 | **63542** | 32260 | **63538** |
| 10% | 32260 | **34944** | 32260 | **34946** |
| 1% | 32260 | **32089** | 32260 | **32086** |
| 0.10% | 32260 | **31803** | 32260 | **31800** |
| 0.01% | 32260 | **31774** | 32260 | **31772** |
| SNP2 (rs10228436) |  |  |  |  |
| Original BAM | **36806** | 34971 | **36806** | 34971 |
| 100% | **73614** | 34971 | **73612** | 34971 |
| 10% | **40531** | 34971 | **40486** | 34971 |
| 1% | **37182** | 34971 | **37174** | 34971 |
| 0.10% | **36843** | 34971 | **36842** | 34971 |
| 0.01% | **36810** | 34971 | **36809** | 34971 |
| SNP3 (rs17337023) |  |  |  |  |
| Original BAM | **29527** | 28865 | **29527** | 28865 |
| 100% | **59057** | 28865 | **59054** | 28865 |
| 10% | **32413** | 28865 | **32479** | 28865 |
| 1% | **29834** | 28865 | **29822** | 28865 |
| 0.10% | **29560** | 28865 | **29556** | 28865 |
| 0.01% | **29530** | 28865 | **29529** | 28865 |

